# Structure and evolutionary history of a large family of NLR proteins in the zebrafish

**DOI:** 10.1101/027151

**Authors:** Kerstin Howe, Philipp H. Schiffer, Julia Zielinski, Thomas Wiehe, Gavin K. Laird, John C. Marioni, Onuralp Soylemez, Fyodor Kondrashov, Maria Leptin

## Abstract

Animals and plants have evolved a range of mechanisms for recognizing noxious substances and organisms. A particular challenge, most successfully met by the adaptive immune system in vertebrates, is the specific recognition of potential pathogens, which themselves evolve to escape recognition. A variety of genomic and evolutionary mechanisms shape large families of proteins dedicated to detecting pathogens and create the diversity of binding sites needed for epitope recognition. One family involved in innate immunity are the NACHT-domain-and Leucine-Rich-Repeat-containing (NLR) proteins. Mammals have a small number of NLR proteins, which are involved in first-line immune defense and recognize several conserved molecular patterns. However, there is no evidence that they cover a wider spectrum of differential pathogenic epitopes. In other species, mostly those without adaptive immune systems, NLRs have expanded into very large families. A family of nearly 400 NLR proteins is encoded in the zebrafish genome. They are subdivided into four groups defined by their NACHT and effector domains, with a characteristic overall structure that arose in fishes from a fusion of the NLR domains with a domain used for immune recognition, the B30.2 domain. The majority of the genes are located on one chromosome arm, interspersed with other large multi-gene families, including a new family encoding proteins with multiple tandem arrays of Zinc fingers. This chromosome arm may be a hot spot for evolutionary change in the zebrafish genome. NLR genes not on this chromosome tend to be located near chromosomal ends.

Extensive duplication, loss of genes and domains, exon shuffling and gene conversion acting differentially on the NACHT and B30.2 domains have shaped the family. Its four groups, which are conserved across the fishes, are homogenised within each group by gene conversion, while the B30.2 domain is subject to gene conversion across the groups. Evidence of positive selection on diversifying mutations in the B30.2 domain, probably driven by pathogen interactions, indicates that this domain rather than the LRRs acts as a recognition domain. The NLR-B30.2 proteins represent a new family with diversity in the specific recognition module that is present in fishes in spite of the parallel existence of an adaptive immune system.

## INTRODUCTION

The need to adapt to new environments is a strong driving force for diversification during evolution. In particular, pathogens, with their immense diversity and their ability to subvert host defense mechanisms, force organisms to develop ways to recognize them and keep them in check. The diversity and adaptability of pathogen recognition systems rely on a range of genetic mechanisms, from somatic recombination, hypermutation and exon shuffling, to gene conversion and gene duplication to generate the necessary spectrum of molecules.

The family of NACHT-domain (Koonin and Aravind 2000) and Leucine Rich Repeat containing (NLR) proteins (reviewed in Proell et al., 2008, Ting et al., 2008) act as sensors for sterile and pathogen-associated stress signals in all multicellular organisms. In vertebrates, a set of seven conserved NLR proteins are shared across a wide range of species. These are the sensor for apoptotic signals, APAF1, the transcriptional regulator CIITA, the inflammasome and nodosome proteins NOD1, NOD2, NOD3/NlrC3, Nod9/Nlrx1 and the as yet functionally uncharacterized NachtP1 or NWD1 (Stein et al. 2007; Kufer and Sansonetti 2011). Other NLR proteins, which must have evolved independently of the conserved NLR proteins, are shared by only a few species, or are unique to a species. Some non-vertebrates, such as sea urchins or corals, have very large families of NLR-encoding genes (Bonardi et al. 2012), but an extreme example of species-specific expansion can be found in zebrafish (Stein et al. 2007). Such species-specific gene family expansions suggest adaptive genome evolution in response to specific environments, most probably different pathogens (Liu et al. 2013).

The zebrafish has become a widely used model system for the study of disease and immunity (Rowe, Withey, and Neely 2014; Goody, Sullivan, and Kim 2014), and a good understanding of its immune repertoire is necessary for the interpretation of experimental results, for example in genetic screens or in drug screens. In a previous study we discovered more than 200 NLR-protein encoding genes (Stein et al. 2007). The initial description and subsequent analyses (Laing et al. 2008; van der Aa et al. 2009) have led to the following conclusions: The zebrafish specific NLRs have a well-conserved NACHT domain (PF05729), with a ∼70 amino-acid extension upstream of the NACHT domain, the Fisna-domain (PF14484, see this paper). This domain characterises this class of NLR proteins and is found in all sequenced teleost fish genomes, but not outside the fishes (Stein et al. 2007). The NLR proteins can be divided into four groups, each defined by sequence similarity in the NACHT and Fisna domain, and these groups also differ in their N-terminal motifs. Groups 1 and 2 have death-fold domains; groups 2 to 4 contain repeats of a peptide motif that is only found in this type of NLR protein (Fig. 1). In the initial description, all of the novel NLR proteins ended with the Leucine-Rich-Repeats, but it was later found (van der Aa et al. 2009) that several of them had an additional domain at the C-terminus, an SPRY/B30.2 domain (PF00622). This domain also occurs in another multi-gene family implicated in innate immunity, the fintrims (van der Aa et al. 2009).

**Fig. 1:**
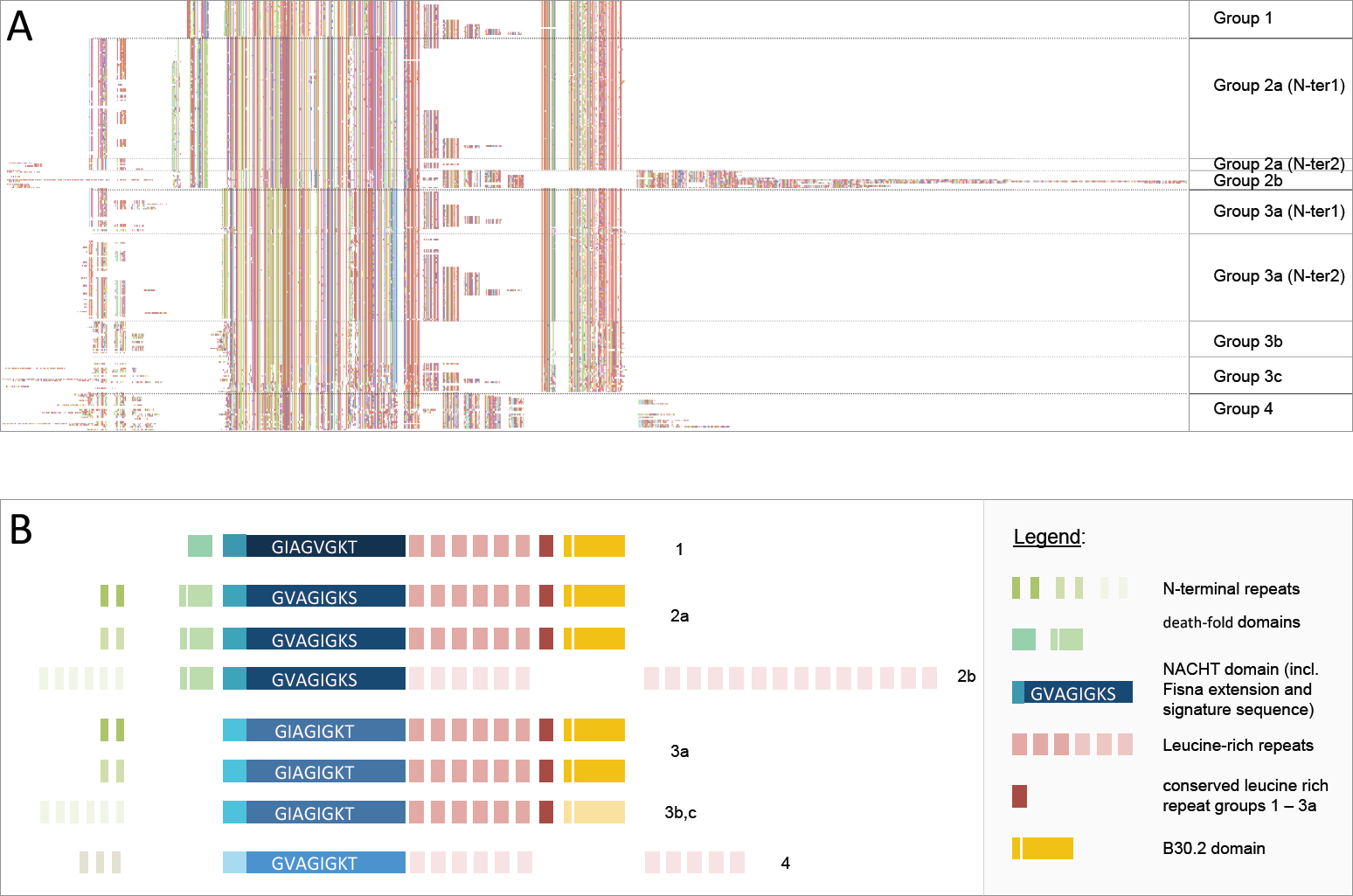
Structure of the fish-specific NLR-B30.2 protein family members. A. Alignment overview of 288 proteins for which full length predictions are available. Regions with long insertions in a few of the genes that had resulted in the introduction of long gaps in the alignment were deleted for the sake of clarity and simplicity. Gaps were manually introduced to highlight intron/exon boundaries, except between the C-terminal extensively duplicated LRRs in group 2b and the extensively duplicated N-terminal repeats in groups 2b and 3b. The colour coding is a random assignment of colours to amino acid created in Jalview. A gap was inserted between the LRRs 6 and 7 of groups 2b and 3b to allow the conserved C-terminal LRR and the B30.2 domains of groups 1, 2a and 3a to be positioned immediately after the 1-6 LRRs in these groups. An alignment of the full set of predictions is provided in the online supplementary material. B. A schematic representation of the protein domains in each group, on the same scale as in the alignment above. Each box represents an exon. Please refer to Supplementary Fig. 2 and the alignment file (online supplementary material) for the many details not captured in this simplified Fig.

The initial identification of the genes and subsequent analyses by others (Laing et al. 2008) suffered from the limitations of the then available Zv6 assembly and gene annotations (published in 2006). In particular, there was a limited amount of data available for long-range assembly arrangements and a lack of supporting evidence for gene models, such as well annotated homologues from other species. In addition, the very high similarity of the NLR genes, as well as their clustered arrangement in the genome further complicated the assembly. As a result, many genomic regions were collapsed and many of the gene models were incomplete. We have revisited this gene family to improve the genome assembly in the regions of interest, have manually re-annotated and refined the gene structures and here provide a full description and analysis. Our re-annotation and analysis of the NLR gene family in zebrafish, and comparison with other species, shows a structured set of more than 400 genes. Different evolutionary pressures on the different parts of the proteins may have created functional diversification between the groups.

## RESULTS

### 1. Identification of all NLR genes in the zebrafish genome

To identify the entire set of fish-specific NLR encoding genes in the zebrafish genome, we used various approaches to collect lists of candidate genes based on the Zv9 assembly (GCA_000002035.2), (for details see Methods and Supplementary Methods). We identified genomic regions containing domain motifs via hmmsearch (hmmer.janelia.org/search/hmmsearch), electronic PCR (Schuler 1997), TBLASTN searches, and by mining the existing annotations for keywords. This collection was purged of gene models belonging to other known families, e.g. fintrims. We identified all overlapping Ensembl and VEGA gene models for the remaining regions of interest. The VEGA gene models were refined and extended through manual annotation and both gene sets merged, resulting in 421 NLR gene models.

Beyond the seven conserved NLR genes, and nine other NLR genes (Table 1) that had a different structure from those described previously and below, the zebrafish genome contains 405 genes encoding NLR proteins that are members of the family we had previously called ‘novel fish NLR proteins‗ (Stein et al. 2007). Henceforth, we will refer to these as NLR-B30.2 proteins (see below). From the 405 genes 37 were pseudogenes leaving 368 predicted protein-coding NLR-B30.2 genes (online supplementary material, Figshare: http://dx.doi.org/10.6084/m9.figshare.1473092). The genome assembly components carrying these gene models were checked for correct placement and relocated if necessary, thereby contributing to improvements for the GRCz10 assembly (GCA_000002035.3).

**Table 1:**
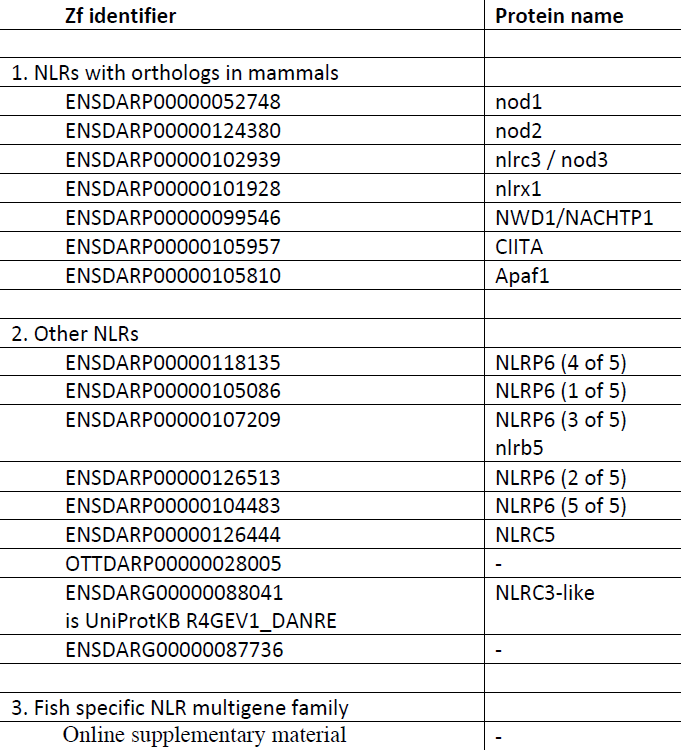
All NLR proteins in the zebrafish genome.

### 2. Structure, conservation and divergence

#### 2a. Domain structure of the NLR family members

The original set of 205 genes described in (Stein et al. 2007) was divided into four groups based on sequence similarity in the Fisna and NACHT domains, and the sequence elements upstream of the Fisna domain. The current dataset shows that all four groups share the Fisna domain, the NACHT domain and the LRRs. Other domains are present only in subsets of the genes; these include a death-fold domain, a B30.2 domain and various N-terminal peptide repeats (Fig. 1 and Supplementary Fig. 1).

*Fisna and NACHT.* The sequence similarities in the Fisna and the NACHT domain in our updated gene set confirm our previous subdivision of the genes into four main groups. A defining motif for each group is the sequence of the Walker A motif, for which the consensus over the whole set is G[IV]AG[IV]AGK[TS] with each group having its own, conserved motif (Fig. 1, Supplementary Fig. 2). Some of the groups can be subdivided further. For example, group 2 consists of two subgroups (Fig. 1, Supplementary Fig. 1), one large and very homogeneous (group 2a), the other smaller, and also homogeneous, with all genes located in a cluster on chromosome 22 (group 2b). Moreover, group 3 has several subgroups that differ in their N-terminal peptides, or their LRRs, or their B30.2 domains (Fig 1.).

We previously found good matches for the Fisna-domain only in fish NLR proteins. Thus, the presence of this domain in conjunction with the particular Walker A motifs is a diagnostic property of the protein family, although short stretches of this domain resembled peptides within mammalian Nod1 and Nod2 (Stein et al. 2007). We used our new collection of Fisna sequences to build an HMM defining a Pfam family (deposited as PF14484) and to search for homologies within mammalian proteins. This revealed alignments with high significance to mammalian members of the NLR family (NLRP3 and NLRP12), with the matching sequence located immediately upstream of the NACHT domain. Secondary structure predictions based on two representatives from zebrafish and the rat using the PSIPRED workbench suggest that the zebrafish Fisna domain may take on the same conformation as the corresponding mammalian proteins (Fig. 2). Together, these findings suggest that contrary to previous assumptions, the Fisna domain was present in the common ancestor of mammalian and fish NLR proteins. Whether it forms a separate domain, or is simply an N-terminal extension of the NACHT-domain, similar to the additional helices seen in the structure of another NLR protein, Apaf1, remains to be determined by experimental work. We did not find Fisna domains in non-vertebrate genomes.

**Fig. 2:**
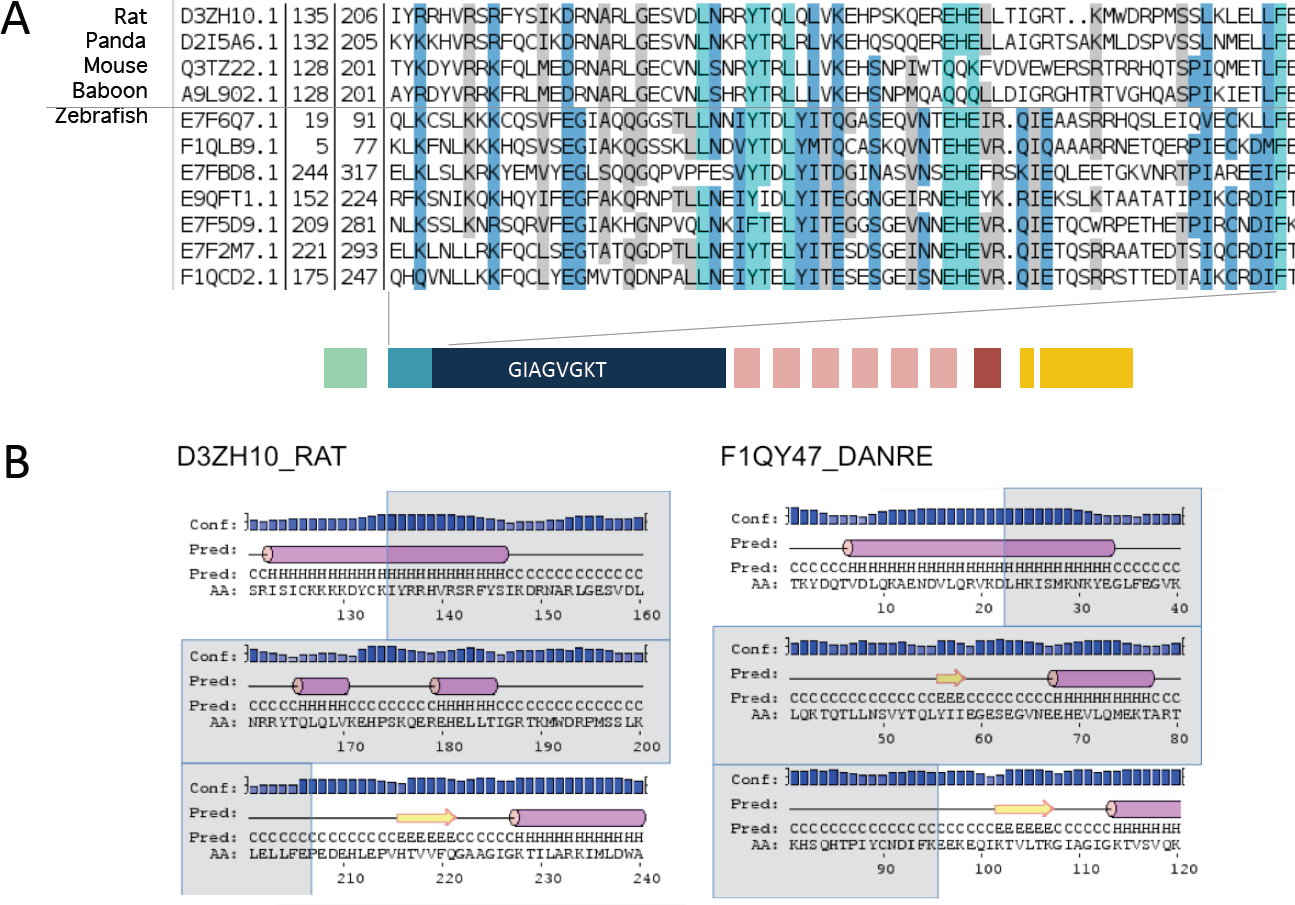
The Fisna domain and its homologs in mammalian proteins. A. Alignment of sequences identified in mammalian genomes using an HMM search with PF14484 and examples of zebrafish Fisna domains. B. Secondary structure predictions for selected examples.

*Death-fold domains and N-terminal repeats.* The four group-specific similarities continue upstream of the Fisna-domain: Groups 1 and 2 both have a death-fold domain; a Pyrin-domain (PYD) with an N-terminal peptide characteristic of BIR domains (http://elm.eu.org/elmPages/LIG_BIR_II_1.html) in the former, and a PYD-like domain in the latter.

The predicted proteins of groups 2, 3 and 4 contain several repeats of a ∼30 amino acid peptide motif. The repeats are not all identical, but occur in two main types. Surprisingly, both of the main types of N-terminal repeat can associate with either group 2 or group 3 NACHT domains (Supplementary Fig. 1: group 2a (N-ter1) and group 2a (N-ter2))., indicating extensive exon-shuffling between family members.

*LRRs.* The proteins of all groups contain several Leucine Rich Repeats (LRRs; Pfam Clan: CL0022). The LRRs in groups 1, 2a and 3 are very similar to each other and occur in a similar pattern; each protein contains between 2 and 7 repeats, with each repeat consisting of two LRRs (Supplementary Fig. 3). There are two types of LRR repeat: the last LRR, immediately upstream of the B30.2 domain, occurs in each gene exactly once, and barely differs between the genes. The other type of LRR in groups 1, 2a and 3 can vary in number between 0 and 6, and are more divergent in sequence. Thus, similar to the situation with the LRRs in the lamprey VLR proteins (Rast and Buckley 2013), the C-terminal LRR seems to be fixed, whereas the others vary more and are duplicated to varying degrees. Groups 2b and 4 do not show this arrangement, but they have yet another type of LRR, which can occur up to more than 20 times.

*SPRY/B30.2 domain*. A B30.2 domain has so far been reported to be present in some but not all fish NLR proteins (van der Aa et al. 2009). Our current set of sequences shows that the presence of the B30.2 domain is restricted to groups 1, 2a and 3, and the domain is missing in groups 2b and 4 (Fig. 1, Supplementary Fig.s 1,4). In view of the extreme similarity between the N-terminal parts (NACHT and death-fold domain) of groups 2a and 2b, and the overall conservation of the gene structure throughout the whole family, it seems most likely that the B30.2 domain was present in the common ancestor of the family, but lost by groups 2b and 4, rather than independently gained by the other groups. We therefore refer to the entire family as the NLR-B30.2 protein family.

#### 2b. Exon-intron structure

All genes in this family have the same exon-intron structure (Supplementary Fig. 2). The largest exon contains the NACHT domain with the N-terminal Fisna extension and the winged helical and superhelical domains, as is also the case in NLRC3, for example. All other domains (N-terminal peptides, LRRS, B30.2 domain) are each encoded on single exons.

#### 2c. Sequence variation between the groups

Visual inspection of the aligned amino acid sequences of all four groups showed that some parts of the proteins were highly conserved, while others showed many amino acid substitutions. The NACHT domains of groups 1, 2a and 3a show little sequence divergence within each group, but strong divergence between the groups (Supplementary Fig. 2). In contrast, there is no recognizable group-specific sequence pattern in the B30.2 domain. Instead, a number of apparently sporadic substitutions are seen throughout the entire set (Fig. 1). This observation is supported by independent phylogenetic analyses of the domains that show different tree topologies (Online supplementary material). The NACHT domains cluster into monophyletic groups, as expected, since similarity in the NACHT domain was used to define the group. In contrast, the tree for the B30.2 domain shows a more interleaved pattern. The discrepancy between the trees suggests different evolutionary trajectories for the B30.2 and NACHT domains. Thus, on the one hand, proteins with different NACHT domains share similar B30.2 domains, and on the other, proteins with nearly identical NACHT domains and N-terminal motifs, such as those in groups 2a and 2b, have different C-termini.

### 3. Evolutionary history

#### 3a. Conservation and divergence within and between the four zebrafish NLR-B30.2 groups

Both the shuffling of N-termini and the unequal divergence of the NACHT and B30.2 domains suggest a complex evolutionary history of the gene family. To analyse the divergence, we calculated rates of non-synonymous and synonymous substitutions in the NACHT and B30.2 domains (dN, dS, and dN/dS values). We studied only those groups that show high intra-group conservation of the NACHT domains (groups 1, 2a, 3a and 3b). We considered groups 3a and 3b separately because inspection of the protein alignment suggested that although they belong into the same group by virtue of their NACHT domain, their B30.2 domains had diverged. We omitted group 3c, as its NACHT domains are more divergent, and may represent further groupings. Median values are in Table 2 and all data displayed in Supplementary Fig. 5.

**Table 2:** Median dN (a), dS (b), and dN/dS (c) values calculated from all pairwise comparisons in the exons coding for the NACHT domain and the B30.2 domain. For dS values below 0.2 are highlighted and for dN/dS values higher than 1 are highlighted.

| **Table 2a** |  | **Fisna-NACHT exon** |  |  |  | **B30.2 exon** |  |  |  |
| --- | --- | --- | --- | --- | --- | --- | --- | --- | --- |
| **dN** |  | Group1 | Group2a | Group3a | Group3b | Group1 | Group2a | Group3a | Group3b |
| Group1 |  | 0.058 | 0.44 | 0.497 | 0.482 | 0.097 | 0.094 | 0.1 | 0.398 |
| Group2a |  |  | 0.029 | 0.411 | 0.398 |  | 0.009 | 0.096 | 0.398 |
| Group3a |  |  |  | 0.038 | 0.11 |  |  | 0.107 | 0.4 |
| Group3b |  |  |  |  | 0.042 |  |  |  | 0.226 |

| **Table 2b** |  | **Fisna-NACHT exon** |  |  |  | **B30.2 exon** |  |  |  |
| --- | --- | --- | --- | --- | --- | --- | --- | --- | --- |
| **dS** |  | Group1 | Group2a | Group3a | Group3b | Group1 | Group2a | Group3a | Group3b |
| Group1 |  | **0.116** | 1.799 | 1.684 | 1.660 | **0.067** | **0.061** | **0.066** | 1.119 |
| Group2a |  |  | **0.043** | 1.525 | 1.491 |  | **0.053** | **0.058** | 1.091 |
| Group3a |  |  |  | **0.038** | 0.216 |  |  | **0.065** | 1.069 |
| Group3b |  |  |  |  | **0.094** |  |  |  | 0.445 |

| **Table 2c** |  | **Fisna-NACHT exon** |  |  |  | **B30.2 exon** |  |  |  |
| --- | --- | --- | --- | --- | --- | --- | --- | --- | --- |
| **dN/dS** |  | Group1 | Group2a | Group3a | Group3b | Group1 | Group2a | Group3a | Group3b |
| Group1 |  | 0.52 | 0.245 | 0.292 | 0.291 | **1.444** | **1.513** | **1.569** | 0.352 |
| Group2a |  |  | 0.639 | 0.267 | 0.267 |  | **1.589** | **1.667** | 0.361 |
| Group3a |  |  |  | 0.942 | 0.506 |  |  | **1.707** | 0.371 |
| Group3b |  |  |  |  | 0.677 |  |  |  | 0.497 |

*Synonymous substitutions.* The median rate of synonymous sequence substitutions in the NACHT domains was very low when comparing members within a group (0.01 - 0.05). The values for comparisons between groups were 20- to 100-fold higher. The only case where the between-group comparison also gave a low value was for group 3a versus 3b, confirming their classification by NACHT domain as one group.

The B30.2 domains showed a different pattern. Similar to the NACHT domain, the median dS values for comparisons within each group ranged between 0.02 and 0.04 for groups 1, 2a and 3a. However, for the B30.2 domain, the values for the between-group comparisons were low: they were minimally or not at all higher than for within-group comparisons. The B30.2 domain sequences of the members of group 3c were more divergent, both from each other those in groups 1, 2a and 3a (Table 2, Supplementary Fig. 5). The patterns of synonymous divergence between groups were clearly different for NACHT and B30.2 domains and confirmed the different behaviour of the two domains we had noted from the alignment of the protein sequences. The observed patterns are most easily explained by gene-conversion (see Discussion).

*Non-synonymous/synonymous substitutions*. Our calculations of the dN/dS values for these comparisons reveal a second evolutionary mechanism acting in the B30.2 domain. We find high median dN/dS values, indicating positive selection acting on the B30.2 domain (Supplementary Fig. 5). Thus, we confirm and extend a previous conclusion for the B30.2 domain in the fintrim proteins (van der Aa et al. 2009), that positive selection might create variation for pathogen recognition in this domain. We discuss below how the combination of positive selection and gene conversion may have created variation of the B30.2 domain throughout the entire family.

#### 3b. Origin of the NLR-B30.2 gene groups

The degree of apparent gene conversion within the NLR-B30.2 gene family makes it difficult or impossible to judge when the groups arose or when they expanded during fish evolution. Moreover, the high divergence between the four groups suggests the split into groups may be old. To explore whether the groups arose in the zebrafish or in an ancestral species, we compared the NLR-B30.2 genes in the zebrafish with those in the closest relative for which a whole genome sequence exists, the carp, as well as NLR-B30.2 genes in other vertebrate genomes (see Materials and Methods).

A tree resulting from a recursive phylogenetic analysis indicates that the split into groups occurred before the zebrafish-carp divergence (Fig. 3). Groups 1, 2, 3a/b and 4 have orthologous relationships between carp and zebrafish: For example, group 2 from zebrafish is most closely related to a group of genes in the carp that is distinct both from other carp NLR-B30.2 genes and from the other zebrafish genes (see also a tree containing all available zebrafish and carp genes in Supplementary Fig. 6).

**Fig. 3:**
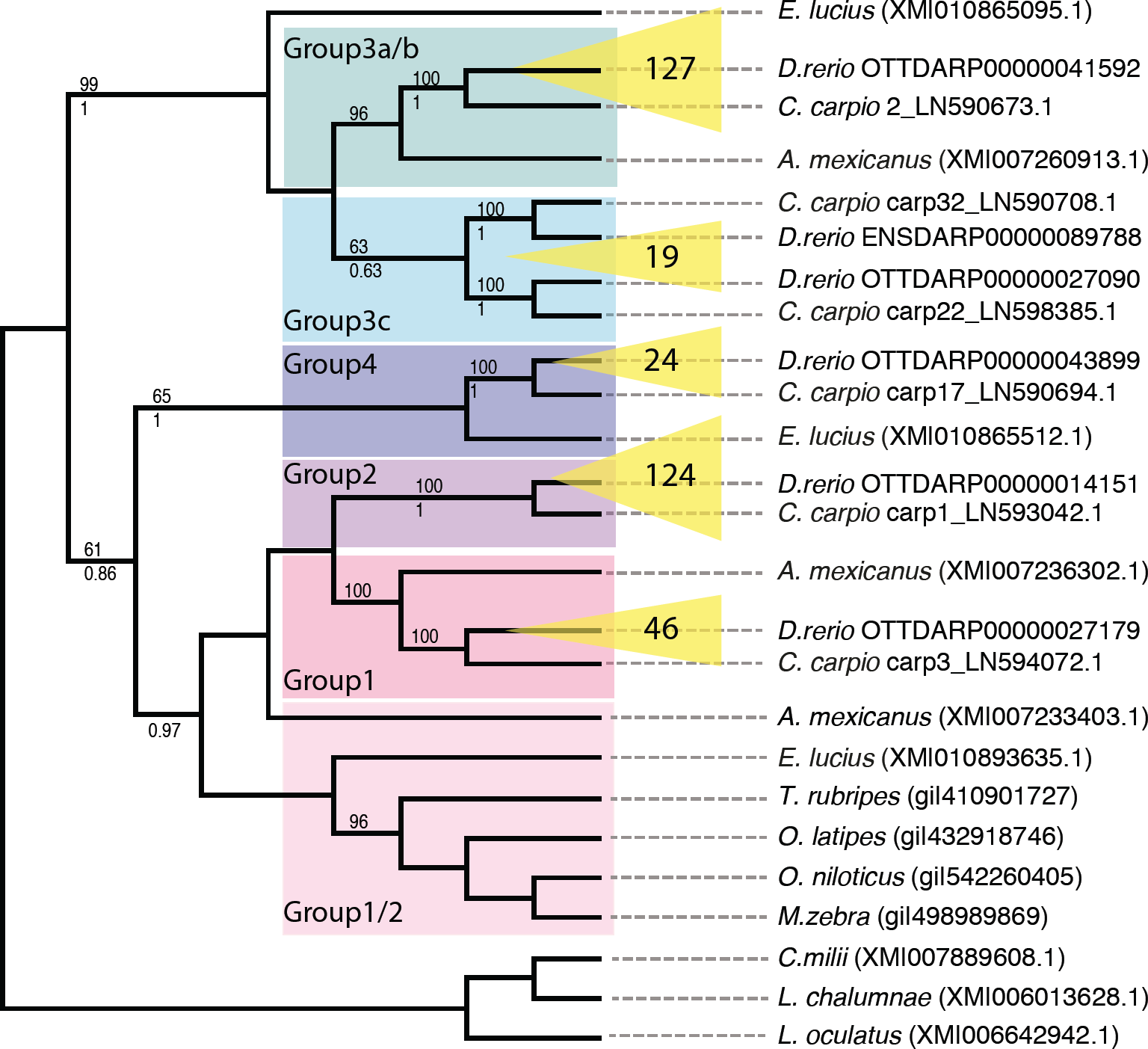
Relationships between NLR-B30.2 genes in different fishes. Phylogenetic tree resulting from a recursive phylogenetic analysis. Inflations of groups in zebrafish are indicated in yellow with numbers of genes per group displayed. Bootstrap values are given above each branch where higher than 50% and pp values (from MrBayes) below each branch, where they are higher than 50% and the topology is congruent.

Not unexpectedly, group 3c, which has a more heterogeneous set of sequences in the zebrafish, shows a more complex evolutionary history. It falls into two groups, both of which have an orthologous group in the carp.

*A mexicanus* and *E. lucius* each have groups of genes that cluster with groups of the zebrafish genes, rather than with each other, but not every group is represented in each of the species. Nevertheless, this suggests that the split is even older than the carp-zebrafish split, having perhaps occurred in basal teleosts. Finally, sequences from *L. oculatus, L. chalumnae* and *C. milii* do not fall into these groups, suggesting that the split into groups must have occurred in the Clupeocephala.

In summary, the groups did not diverge independently from duplicated ancestral genes in each species, but already existed in a common precursor. By contrast, within a species, the majority of genes arose by independent amplification of a founding family member.

#### 3c. Co-occurrence of the NACHT and B30.2 domains

As first reported by van der Aa et al. 2009, the NLR-B30.2 domain fusion is found in all teleost fish. Our collection of NLR-B30.2 genes showed that this domain fusion arose prior to the common ancestor of teleosts, as it was also present in *L. oculatus* (for example ENSLOCG00000000593), for which the genome sequence was not previously available. The NLR-B30.2 proteins predicted in *L. oculatus* also contain an N-terminal Fisna extension, though only distantly related to those in the teleosts, indicating that the ancestral gene included this extension. This is consistent with the fact that the N-terminal extension of the mammalian NLRP3 proteins has recognizable similarity to the Fisna domain, as described above.

We do not find evidence of the NLR-B30.2 fusion in any of the tetrapod genomes, nor in the *L. chalumnae* or *C. milii* genomes. These results indicate that the fusion occurred at least in the common ancestor of the Neopterygii, prior to the third whole genome duplication in the teleost lineage. Genome sequence data from sturgeon, paddlefish or bichir clades, which are currently not available, would provide further information on the point of emergence of the NLR-B30.2 fusion.

#### 3d. Expansion of the NLR-B30.2 family in fish

The phylogeny of the NLR-B30.2 gene family also provides information on the timing of the lineage expansion of the NLR-B30.2 genes observed in the zebrafish. The *C. carpio* genome contains a similarly large family of NLR-B30.2 proteins, but due to the polyploidy of the species some of the nearly identical sequences may constitute alleles rather than paralogues. *A. mexicanus*, a direct outgroup to the zebrafish-carp clade, features the second largest NLR-B30.2 gene family with ∼100 members, showing that the lineage expansion began prior to the zebrafish-carp split. Other fish species, including the spotted gar have fewer than 10 NLR-B30.2 genes, while *E. lucius*, a Euteleost, has ∼50. Thus, the initial gene expansion either occurred in the basal branches of Teleostei with a subsequent loss in some lineages, or independently in several lineages. Independent expansions and losses are a likely scenario, given the expansions of NLR genes in many other species, such as sponges and sea urchins (Yuen, Bayes, and Degnan 2014; Lapraz et al. 2006). The results on fish show that expansions of the NLR-B30.2 family genes began as soon as the NLR-B30.2 fusion occurred, with different dynamics in different lineages.

#### 3e. Age of the NLR-B30.2 family relative to conserved NLRs

We compared the origin of the NLR-B30.2 gene family to the evolutionary history of other NLR genes. There are many species-specific expansions of NLRs, such as the Nalp proteins in mouse and human and the NLR-B30.2 genes in fish, as well as independent inflations in Amphioxus, sea urchin, and sponge. However, there are also the seven NLR proteins that are conserved in all vertebrates and show orthologous relationships. We collected the available orthologues of the conserved NLRs from key metazoan species and created an alignment. We also included all NLR genes from fish that did not belong to the NLR-B30.2 group (listed above and in M&M).

We found that two of the conserved vertebrate NLR genes appear to be shared by all animals (Fig. 5, Supplementary Fig. 6, and online supplementary material). The genes for NWD1 (first described in zebrafish as NACHT-P1) and Apaf1, must have been present in the last common ancestor of bilaterians and non-bilaterians, as they are found in sponges, cnidarians, and all bilaterians analysed. We could not find any candidates in comb jellies (ctenophores). The other five conserved NLR proteins - Nod1, Nod2, NLR3C, CIITA and NLRX1 – arose later in evolution, at the base of the gnathostomes. An additional gene, NLR3c-like, was present at this point, but appears to have been lost in the tetrapod lineage.

In summary, all of the conserved vertebrate NLRs are older than the NLR-B30.2 family. They are never duplicated and certainly not expanded to higher gene numbers in any species.

Important events in the evolutionary history of the NLR-B30.2 family and conserved NLR proteins are summarized in Fig. 4 and Supplementary Fig. 7. The oldest metazoan NLR proteins are NWD1 and Apaf1, while Nod1, Nod2, NLR3C, CIITA and NLRX1 and NLR3C-like were acquired in the lineage leading to gnathostomes. A fusion of the NLR and B30.2 domains then occurred in the Neopterygian lineage and subsequently these genes diversified into groups, within which they continued to expand in various fish lineages. The high similarity of the NACHT domains within the groups appears to have been maintained by gene conversion. Large-scale amplifications of NLR genes occurred independently in many species.

**Fig. 4:**
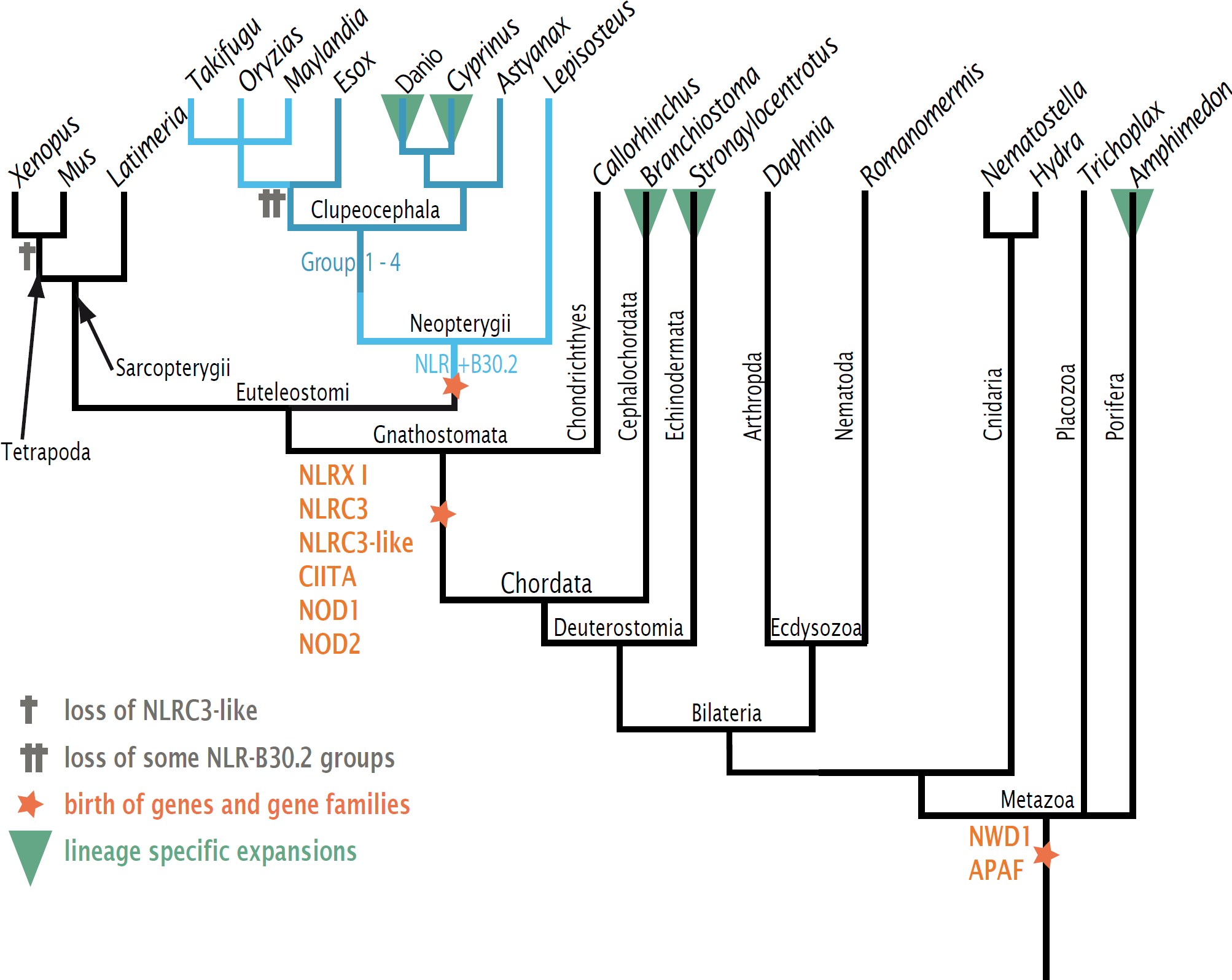
Evolutionary history of NLR genes A reduced dendrogram of Metazoa based on the NCBI taxonomy database displaying key events in the evolution of NLR genes as described in the main text. See Supplementary Fig. 7 and the supplementary online material for phylograms.

### 4. Genomic location

The first survey of NLR genes on the zebrafish genome assembly Zv6 suggested that they were located on 22 different chromosomes, with some enrichment on chromosome 4 (50 genes) and chromosome 14 (47 genes) (Stein et al. 2007). Since this analysis, the assembly of the zebrafish genome has been significantly improved and the current Zv9 NLR gene set shows a more restricted distribution (Fig. 5), with 159 (44%) of the genes located on the right arm of chromosome 4. The remaining genes are distributed between 12 other chromosomes (153 genes) and unplaced scaffolds (56 genes).

**Fig. 5:**
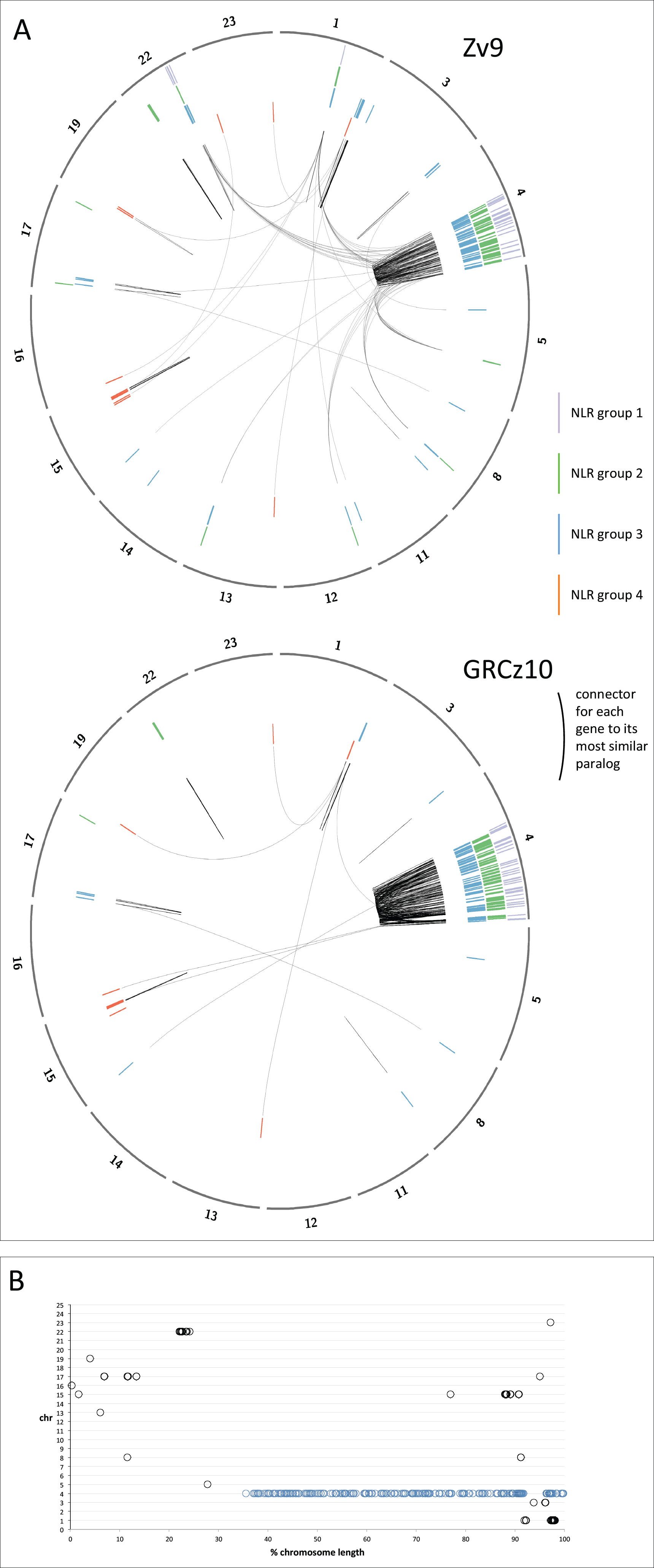
Location of the NLR genes in the genome in assemblies Zv9 and GRCz10. A. The chromosomes containing NLR-B30.2 genes are shown in the outer circle (note that corrections of the genome between Zv9 and the GRCz10 have changed the lengths of some of the chromosomes. The genes were annotated on Zv9 and lifted over to the GRCz10 path where possible as the GRCz10 gene set did not become available until May 2015. The members of the four groups of NLR genes are shown as radial lines within the circles with group 1 in the outermost and group 4 in the inner ring. Each gene is connected by a black line to its most closely related paralog, based on the number of amino acid substitutions per site calculated in MEGA5 (Poisson correction model). Most genes are most closely related to a near neighbour. The changes in the assembly have have lead to many genes that were closely related but resided on different chromosomes in Zv9 being located in closer proximity in GRCz10. B. Normalised location of NLR-B30.2 genes on chromosomes. Each chromosome is shown as a horizontal line of 100% length, and the NLR genes are plotted at their relative positions along the chromosome. Apart from the genes on chromosome 4 (marked in blue), all other genes are found within the first or last quarter of the chromosome.

Additional sequencing and data gathering by the Genome Reference Consortium since the release of Zv9 led to the rearrangement of multiple assembly components, including relocation of sequence to different chromosomes. These placements are based on manual curation by the Genome Reference Consortium, supported by genetic mapping data, clone end sequence placements and optical mapping data (Howe and Wood 2015). The latest assembly, GRCz10, reveals that the majority of the genes clustered on the long arm of chromosome 4, where 75% of the NLR-B30.2 genes, including all group 1 and group 2a genes, now reside (Fig. 5). Group 2b genes are now found exclusively in a cluster on chromosome 22, which suggests that they arose via local duplications of a single precursor gene that had lost its B30.2 domain. Similarly, group 3a genes are clustered together on chromosome 4, with group 3b and 3c genes clustered on chromosomes 1 and 17, respectively. Group 4 genes are found mostly on chromosome 15, some on chromosome 1, with none found on chromosome 4. Both group 2 and group 3 have a few individual genes dispersed over other chromosomes; careful inspection of the evidence on which this allocation is based revealed no indications that it is incorrect. Some of the group 3 members on other chromosomes are more divergent from the consensus for this group, suggesting they may indeed have separated from the group early.

Within chromosome 4, no clear pattern can be detected in the distribution of the genes. We are, however, aware of possible shortcomings in the assembly of the long arm of chromosome 4; the highly repetitive nature of the sequence makes it difficult to exclude with absolute certainty shuffling of gene locations. In addition to containing multiple copies of 5S ribosomal DNA (Sola and Gornung 2001), 53% of all snRNAs and the majority of the NLR-B30.2 genes, chromosome 4 also contains multiple copies of genes encoding a particular type of Zn-finger protein, which we discuss below.

Finally, another striking feature of the genes’ genomic location is that they tend to accumulate near the ends of the chromosomes (Fig. 5b). With the exception of the cluster on chromosome 22, and two single genes on Chromosomes 5 and 15, all other genes (81% of the NLR genes outside chromosome 4) are located within 15% of chromosome ends. On chromosome 4 we found 26% of the genes within 15% of the end.

### 5. Distribution of Fintrim and multiple Zn-finger encoding genes

We noticed that the NLR-B30.2 genes on chromosome 4 were often interspersed with genes encoding multiple tandem Zn-finger proteins. In some cases, gene models had been made that joined B30.2 domains with Zn-fingers, but our analyses showed that the B30.2-encoding exons instead belonged to a neighbouring NLR gene, rather than the more distant Zn-finger encoding exons. A possible explanation for the mis-annotation is that the predictions created apparent Fintrim genes. Fintrim proteins are composed of multiple Zn-fingers combined with a B30.2 domain and are assumed to act as sensors for immune stimuli (van der Aa et al. 2012). They are often found in the vicinity of, or at least on the same chromosome as, genes that are located in the MHC in mammals (MHC-associated genes are found not in one cluster in the zebrafish, but on four different chromosomes [3, 8, 16, 19]). We therefore analysed the distribution of the NLR-B30.2 genes relative to the location of fintrims and multiple Zn-finger encoding genes.

To establish a list of fintrim genes from the current assembly, we collected and refined candidate genes from the Zv9 genes set, resulting in 61 TRIM, 40 ftr and 18 btr genes (online supplementary material). An alignment of 283 B30.2 domains from the NLR-B30.2 genes and 117 from the fintrim and btr families shows that most of the B30.2 domains from the NLR-B30.2 genes are more closely related to each other than to those of the TRIM families. We found no close association in the genome between the fintrim and the NLR-B30.2 genes (Fig. 6B), except for two cases where a single fintrim gene is found near an NLR-B30.2 gene cluster (chr1 and chr15). If anything, fintrim genes are excluded from regions where NLR-B30.2 genes are found and vice versa.

**Fig. 6:**
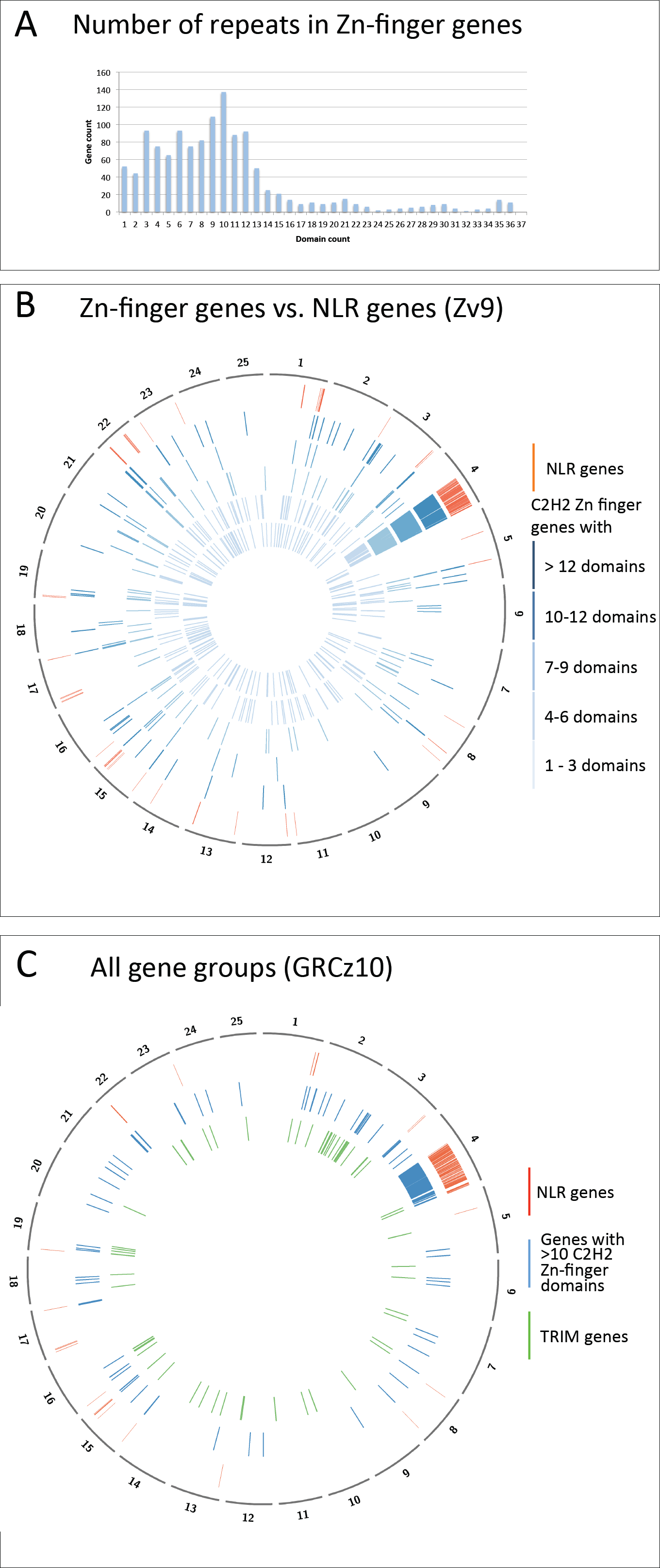
Genomic positions of genes for multi-Zn-finger proteins, Fintrims and NLR-B30.2 proteins. A. Frequency of genes encoding proteins with the indicated number of Zn-finger domains. Total number of Zn-finger encoding gene predictions: 1102. B. The genomic locations of Zn-finger encoding genes is plotted on a circos diagram representing the Zv9 assembly for five subgroups defined by the number of Zn-finger domains. Inner circles (light to dark blue): 1-3, 4-5, 6-7, 8-9, 10-12 and more than 12 domains; Outer circle (red): NLR genes C. Spatial distribution of NLR genes, TRIM genes and genes with at least 10 Zn-finger domains in the GRCz10 assembly (for gene liftover see Fig. 5).

However, as noted above, genes encoding multiple Zn-fingers were interspersed among the NLR-B30.2 genes. Unlike the TRIMs, which contain C3HC4 (RING) and Znf-B-box domains (IPR001841 and IPR000315, respectively), the genes on chromosome 4 encoded yet uncharacterised proteins consisting exclusively of tandem repeats of Zn-fingers of the classical C2H2 type (IPR007087). To investigate this new gene family further, we collected all gene models encoding Zn-fingers of this type. A total of 1259 gene models were found, with the number of repeats per gene ranging from 1 to 36. Some of the cases of genes with extremely large numbers of C2H2 domains (e.g. ENSDARP00000109314) may be mis-annotations that combine adjacent, shorter genes, which we did not analyse manually in further detail.

The encoded proteins with small numbers of Zn-fingers included many known proteins, including the Sna and Opa transcription factor families. These were broadly distributed in the genome and largely excluded from chromosome 4 (Fig. 6). By contrast, those with larger numbers of Zn-finger domains are progressively clustered in restricted regions of the genome. For example the majority (66%) of genes with more than 10 C2H2 domains are found on the right arm of chromosome 4, where they are interspersed in an irregular pattern among the NLR-B30.2 genes. Outside chromosome 4, some multi-Zn-finger genes co-locate with subsets of the TRIM genes, for example on chromosomes 3, 16 and 19, while others are located in regions where neither NLR-B30.2 genes nor TRIMs are found. Similar to the NLR-B30.2 genes, multi-Zn-finger genes outside of chromosome 4 tend to be close to chromosomal ends (62% of genes within 15%). On chromosome 4, 8% of the multi-Zn-finger genes are found within 15% from the chromosome ends.

In summary, the local duplications that may have led to the expansion of the NLR-B30.2 genes on chromosome 4 may also have duplicated the multi-Zn-finger genes, which have subsequently been transposed to other chromosomes.

## DISCUSSION

### Phylogeny of vertebrate NLR proteins

The family of NLR-B30.2 genes has been shaped by different genomic and genetic mechanisms throughout evolution. These include repeated gene amplifications, shuffling of exons and gene fusions, gene conversion and positive selection for diversity. We see low rates of synonymous substitutions in the NACHT domain when comparing the members within each group, and similarly low rates of synonymous substitutions for the B30.2 domains in comparisons across all groups. A low rate of synonymous sequence substitutions can be interpreted as a sign of recent gene duplication. If we apply this interpretation using the low divergence of the B30.2 domain, then we would have to conclude that the entire set of genes in groups 1, 2a and 3a is the product of recent duplications. However, this is not consistent with the significant divergence of the NACHT domains between the groups, the different tree topologies of the two domains, nor with our finding that the split into NLR families occurred before the divergence of the zebrafish and the carp. Therefore there must be an alternative explanation. The pattern of synonymous divergence of the two domains between groups is most parsimoniously explained by ongoing gene conversion in the NACHT-B30.2 gene family, with conversion in the NACHT domain acting only within each group, whereas the B30.2 domains show signs of gene conversion across the whole set of genes of groups 1, 2a and 3a, i.e. also between members of different groups.

Gene conversion is not uncommon in gene families involved in immunity (see Pasquier 2006 for review). It can create diversity, for example in antibodies (Wysocki and Gefter 1989) or in the MHC (reviewed in Martinsohn et al., 1999), but it can also homogenize genes, e.g. in the T cell receptor family (Jouvin-Marche, Heller, and Rudikoff 1986). In the NLR-B30.2 both mechanisms may operate.

Gene conversion in the NACHT domain appears to be restricted to conversion within groups, keeping the groups (a) homogeneous and (b) distinct from each other. In contrast, gene conversion between B30.2 domains may have another effect, namely to create additional variation. The high dN/dS values indicate positive selection for non-synonymous variants in residues potentially involved in pathogen recognition. As noted by (van der Aa et al. 2009) for fintrims, and also seen in the NLR-B30.2 proteins, the substitutions are concentrated in regions of the B30.2 domain that are exposed on the surface and likely involved in pathogen interactions. Once substitutions have been introduced in one of the genes, gene conversion can then spread these throughout the family. If conversion is frequent and the conversion tracts are short, i.e. cover only a small part of the B30.2 exon, then it would recombine individual amino acid replacements occurring in different parts of the domains and in different paralogues. The process would create novel combinations of amino acid substitutions, and these variants would then again form the substrate for further selection. At the same time, since gene conversion in the B30.2 domain acts across groups, this mechanism also ensures that new recognition modules can spread beyond the group in which they first arose. This can prevent the groups being locked in on a defined subset of B30.2 domains. It is striking that the three groups of genes that show gene conversion in the B30.2 domain are all localised on chromosome 4, whereas group 3b, which has diverged from group 3a in its B30.2 domain, is located on chromosome 1.

The oldest NLR genes appear to be those encoding the ancestors of two conserved NLRs, Apaf1 and NWD1, which we find in all animal lineages. These proteins have not been reported to have immune functions. Apaf1, originally discovered as CED-4 in *C. elegans*, is an ancient regulator of apoptosis. The so far only function reported for NWD1, first identified in the zebrafish genome as NACHTP1 (Stein et al. 2007), is its involvement in androgen signalling in the context of prostate cancer (Correa et al. 2014). It will be interesting to learn whether this is a special case of a more general immune function yet to be discovered, or whether, like Apaf1, this old gene does not have immune functions. The other conserved genes first appear at the base of the gnathostomes, and all have roles in immunity or inflammation - whether as transcription factors or as inflammasome components.

In parallel, NLR genes have duplicated and often undergone extensive species-specific expansions throughout evolution. This is the case, for example, for the members of the Nalp/NLRP family in the mouse and the NLR-B30.2 family we discuss here. The largest of the known early expansions were in the sponge *A. queenslandica*, the sea urchin *S. purpuratus* and the lancelet *B. floridae*, with about 120, 92, and 118 genes respectively (Yuen, Bayes, and Degnan 2014; Huang et al. 2008). As more genomes are sequenced it is likely that additional NLR expansions will be discovered. In vertebrates, the largest expansions are those of the NLR-B30.2 family, although we also find other NLR gene families, for example in the elephant shark *C. milii* (Supplementary Fig. 7, online supplementary material).

The expansion of the families argues in favour of their involvement in immunity or broader stress reactions, as seen in numerous other examples of expanded gene families. Expansions can increase the amount of gene product, for example to adapt to stressful environmental conditions (Kondrashov et al. 2002; Kondrashov 2012), as in the cold adaptation in several gene families expressed in Antarctic notothenioid fishes (Chen et al. 2008). Expansions can also allow the creation of the variety of sequences that is needed for immune recognition, as in the case of antibodies and T-cell receptors, or the more recent example of the VLR genes in lampreys and hagfish (Li et al. 2013).

One possible scenario for the creation of the current NLR-B30.2 gene family in the zebrafish is as follows. Early in the fish lineage, after their initial creation through the fusion of the NLR and B30.2 components, the NLR-B30.2 genes underwent duplications, similar to many other NLR genes in other lineages (Hamada et al. 2012; Yuen, Bayes, and Degnan 2014). At this point, the available data do not allow us to trace these earliest duplications.

In the Clupeocephala, the paralogues then diversified into groups. Whether the common ancestor had four genes (or a similarly small number), or whether each of the four genes had already begun to be duplicated to form small families is not clear. At this point, gene conversion may already have been occurring and if the early prototypes had already amplified into gene families in the common ancestor, then gene conversion may have acted within each group. What seems clear is that if gene conversion ever acted across the whole gene, conversion of the NACHT domain between groups must have stopped when the groups became too divergent. It cannot have acted between the NACHT domain encoding exons of different groups, as the differences between the groups have been maintained and are still visible now. Since not all currently extant fish have representatives of all four groups, it may be that either whole sets of these genes can be easily lost, or else that the common precursor had only one gene from each group, and that not all lineages inherited all four prototypes. The near-identity of some of the genes we find in the zebrafish (difference between paralogues lower than rate of polymorphism) shows that duplications continue to occur.

It is worth speculating about the functional and selective forces that prevent sequence homogenization between the NACHT domains of different groups. If the proteins form large multimeric complexes, as the known inflammasome NLRs do, then their efficient functioning might require that only proteins from the same group can multimerise, for instance to elicit distinct down-stream signaling events. This is supported by the groups featuring different N-terminal domains. A mixed multimer may not be able to assemble a functional N-terminal effector complex.

The C-terminal domains - LRRs and B30.2 - do not show the same clear subdivision into families as the N-terminal and the NACHT domains, and homogenizing gene conversion must therefore have affected only part of each gene, or affected different parts differently. This is not without precedent, since gene conversion often proceeds across DNA segments of limited length (see Innan 2009 for review) and parts of a gene can escape sequence homogenization (Innan 2009; Teshima 2004).

Both LRRs and B30.2 domains have been implicated in recognition of pathogen- or danger-associated molecular patterns. The B30.2 domain of Trim5a binds to HIV-1 and is involved in blocking HIV-1 proliferation in monkeys (Stremlau et al. 2004). In Nod1 and Nod2, the LRRs recognise components of pathogens (Inohara et al. 2005), including flagellin monomers (Akira, Uematsu, and Takeuchi 2006), while in Toll-like receptor proteins they recognise viral dsRNA (Kawai and Akira 2009).

The sequences of the LRRs in the NLR-B30.2 genes are not particularly variable, and it therefore seems unlikely that they have a role in specific ligand-recognition. The B30.2 domains however show significant amino acid variation between the members. It may therefore have the same function as the related B30.2 domain in the fintrim genes, which has been suggested to be under positive selection to allow variation in specificity for pathogen recognition. It is conceivable that the acquisition of the B30.2 domain and the option to use it for specific recognition of a wide range of pathogens drove the amplification of these genes.

While we lack sufficient genomic data from salt-water fishes, we are tempted to speculate that the massive inflation of the NLR-B30.2 group may have occurred along with the adaptation to fresh water environments when ancestors of the zebrafish encountered a new pathogen fauna. Alternatively, the NLR-B30.2 system may functionally complement the innate immune system during the first few weeks of life of the zebrafish larva: the larva is exposed to the outside world and starts eating after two days of development, but a functional adaptive immune systems arises only after three to five weeks (Lam et al. 2004). We have not investigated whether the presence of NLR-B30.2 expansions in a fish species correlates with the time of development of the adaptive immune system in that species.

### Shuffling between genes and creation of new genes

A mechanism involved in the initial creation of the NLR-B30.2 family appears to have been exon shuffling, both within the family and between the NLR genes and other gene families. For example, the N-terminal peptide repeats occur in several variants, but a given variant is not strictly associated with any particular group: at least two of the variants are found both in association with group 2a and group 3.

We also find evidence for recombination with other immune genes. The B30.2 domain of the NLR-B30.2 proteins most closely resembles that of the fintrim proteins, a fish-specific gene family for which the origin of the fusion between the Zn-fingers with the B30.2 domain is not known (van der Aa et al. 2009), suggesting that exon shuffling occurred during the generation of the ancestral genes of the NLR-B30.2 and the fintrim gene families.

Apart from this possible case of exon-exchange, the relationship between the three large and partially related families – the NLR-B30.2 genes, the fintrim genes and the multi-Zn-finger genes we describe here – are unclear. While it is striking that the fintrims share the B30.2 domain with the NLR-B30.2 genes and the Zn-fingers with the multi-Zn-finger proteins, they do not preferentially map to the same regions of the genome, and the Zn-finger is of a different type. By contrast, the multi-Zn-finger genes are mostly found on chromosome 4, interspersed between the NLR-B30.2 genes. We have not attempted to trace the relative evolutionary histories of these three families.

A further gene that may have arisen from domain shuffling between these gene families is the human gene encoding pyrin (marenostrin/MEFV). Pyrin is a protein that is composed of an N-terminal PYD domain, for which the best match in the zebrafish is the PYD domain in the group 1 NLR-proteins. The C-terminal part of pyrin contains a Zinc-finger and a B30.2 domain, which resembles the zebrafish fintrim proteins of the btr family. The most likely interpretation for the origin of this gene, which must have arisen at the base of the tetrapods, is therefore a recombination between an NLR gene and a neighbouring fintrim gene.

### Chromosome 4

The zebrafish chromosome 4 has unusual properties. Its long arm is entirely heterochromatic, replicates late and shows a reduced recombination rate. It contains an accumulation of 5S rRNA, snRNA, tRNA and mir-430 clusters (Anderson et al. 2012; Howe et al. 2013), as well as the expanded protein coding gene here.

Chromosome 4 was recently shown to function as the sex chromosome in wild zebrafish ZW/WW sex determination, with the sex determining signal being located near the right end chromosome 4 (Wilson et al. 2014). The sex determination region in the grass carp may also be associated with NACHT domain encoding genes (Y. Wang et al. 2015). This was concluded from the comparison of the genome sequences of one male and one female carp, where those regions present in the male and absent in the female were interpreted as sex-determining. In addition to the NACHT domain genes, this region also included other immunity genes, such as the immunoglobulin V-set, ABC transporters, and proteasome subunits. While the co-location between sex determination and immune signalling molecules we describe here may support this conclusion, it is of course equally possible that the finding in the grass carp is simply caused by allelic diversity in these highly variable genes between the two individuals. It is nevertheless intriguing that two fast evolving genetic systems are located in such close proximity in zebrafish. Perhaps, after an initial round of NLR gene duplications, a run-away evolutionary process of further amplification to created the present chromosome 4, which is now a hotspot for rapid evolutionary processes.

## METHODS

### Re-annotation of NLR genes in the zebrafish genome

To establish a complete list of all genes encoding NLR proteins in the zebrafish genome, we first conducted a search of the Zv9 genome assembly for sequences that encoded the characteristic protein domains, using a combination of approaches. We constructed a hidden Markov model (HMM) for the Fisna domain and used this together with the HMM for the NACHT domain obtained from PFAM to search the Zv9 assembly with hmmsearch (hmmer.janelia.org/search/hmmsearch), resulting in 297 Fisna and 328 NACHT locations (see online supplementary material). As an alternative way to identify NACHT domains specific for the novel NLRs, we ran electronic PCRs (PMID: 9149949) with primer sets for a segment stretching from the C-domain into the winged helix domain that we had used for experimental analysis of the genes (unpublished work). Each set of primers was specific for one of the NLR groups (Supplementary Methods). This resulted in 321 hits. To find regions in the genome encoding B30.2 domains we conducted a TBLASTN search, which yielded 503 hits. As B30.2 domains also occur in other large, immune-related protein families (see below), such a high number of domains was consistent with expectations.

Secondly, we collected all Ensembl genes overlapping the above motifs (487 predicted genes) and also all manually annotated genes (vega.sanger.ac.uk) that had been marked as NLR or as containing a NACHT domain during manual annotations in the past (307 predicted genes).

The collection was purged of gene models that did not match the criteria for being novel NLRs, excluding e.g. the B30.2 domain-containing fintrim genes. Sixteen NACHT domain proteins in the combined list do not belong to the group of novel fish NLRs since they do not contain the Fisna-domain, and the sequence surrounding their WalkerA motifs does not match the one typical for the novel NLRs. They include the seven conserved NLRs that are orthologous across all vertebrate species (Nod1, Nod2, Nlrc3, Nlrx1, CIITA, Apaf1, NWD1/NachtP1), and nine further proteins with an NLR structure. (Table 1 and online supplementary material)

Comparison of the purged gene sets with the genomic regions that encoded parts of NLR proteins showed that many genes in this family had been annotated incorrectly, and for others there were no predictions at all, probably due to the repetitive nature of this gene family and the limited availability of supporting evidence in the form of cDNAs.

The regions containing the sequences identified in our searches were therefore re-annotated manually, correcting and adding gene models to create full-length genes. This re-annotation had to be restricted to regions located on finished sequence, since whole genome shotgun (WGS) contigs in Zv9 were not accessible to manual annotation. For these contigs, the automated Ensembl gene models were retained in their original form, recognizable in our final list by their “ENSDARG” identifier (Supplementary Excel Table1). The resulting protein sequences were then aligned using Clustal-Omega (Sievers et al. 2011) or Muscle (Edgar 2004) and compared. Sequences that appeared truncated were analysed further by searching for additional exons to complete them, until, in an iterative process, we had optimised them. Some sequences remained incomplete, either because they were located next to sequence gaps, or because no additional exons could be detected. In these cases it is not known whether the truncation of the gene is a true biological event caused by recent recombination, or whether it is due to a mis-assembly of the genome sequence.

The optimised gene set was combined with the gene predictions in Ensembl (Methods, hand-filtered alignments and location checks of the remaining genes to identify accordance). The final list of novel zebrafish NLR proteins contains 368 members (Supplementary Excel Table1). A further 36 predictions for NLR genes had been annotated as pseudogenes and were therefore not retrieved for this list (online supplementary material). The refined genes have since been integrated into the VEGA and Ensembl gene sets, however since the annotation was performed on pre-GRCz10 paths, the latest GRCz10 gene set (Ensembl80) might differ marginally from the described results.

### Conserved NLR genes across metazoa

We used the zebrafish gene identifiers for the conserved NLRs in zebrafish to query the Orthoinspector 2.0 database (Linard et al. 2014) at http://lbgi.igbmc.fr/orthoinspector for orthologs in published genomes and downloaded the corresponding sequence. We then queried a custom Blast database of the Cyprinus carpio proteome, as well as the NCBI nr database for selected fish species using BLASTP. After removing redundant hits we calculated alignments employing Clustal-Omega v.1.2 (Sievers et al. 2011) and subsequently removed sequences of poor quality. In a second inference we also used trimal (Capella-Gutiérrez, Silla-Martínez, and Gabaldón 2009) to reduce the alignment to the conserved residues. We employed prottest v.3.2 (Darriba et al. 2011) to infer the best fitting evolutionary model and found that the LG model with Gamma optimisation performed best under the AKIKE information criterion. We then ran RAxML v7.7.2 (Stamatakis 2006) on both alignments on the Cologne University CHEOPS super computer and calculated bootstrap values. Phylogenetic trees were visualised and edited in Dendroscope v.3.2.5 (Huson et al. 2007).

### figmop and tblastn screen for NLR-B30.2 candidates in other fish genomes

Expanded gene families are not well annotated in most genomes. Rather than relying on gene predictions for identifying NLR-B30.2 genes, we therefore directly searched the genome sequences of 6 species: *Latimeria chalumnae, Lepisosteus oculatus, Callorhinchus milii, Esox lucius, Astyanax mexicanus, Cyprinus carpio*. We downloaded genome data either from NCBI servers or the genome project websites. We then used the Figmop (Curran, Gilleard, and Wasmuth 2014) pipeline to find contigs and scaffolds in the genomes with NLR-B30.2 candidates on them. The Figmop pipeline builds a profile of conserved motifs from a starting set of sequences and uses these to search a target database with the MEME software suite (Bailey et al. 2009).

We used zebrafish NLR-B30.2 sequences from all four groups to create a set of 15 motifs to earch the above genomes. The resulting contigs were then subjected to the Augustus (v3.0.3) gene prediction pipeline (Stanke and Waack 2003) to predict genes *de novo*, setting zebrafish as the “species”. We complemented this approach by TBLASTN searches using the NACHT as well as the B30.2 domains as queries in individual searches and then kept those predictions in which the domains occurred in the proper order (thereby excluding spurious cases caused by mis-assembly or incomplete genes).

### Phylogenetic analyses of the NLR-B30.2 groups

We used a recursive approach for identifying genes for the phylogeny that were representative of the overall sequence divergence in the gene family. We selected only those that had both the NACHT and B30.2 domains. We then recursively performed the following: constructed a sequence alignment of ∼500 residues (starting with the NACHT domain) in the dataset using Clustal-Omega, (2) constructed a phylogeny using a Bayesian approach using MrBayes with mcmc=1,000,000, sump burnin=1000 and sumt burnin=1000 and (3) removed monophyletic paralogs from the dataset. The recursive analysis was halted when no instances of paralogous sister-sequences remained (Fig. 3), with the exception that at least one zebrafish and carp sequence from each of the major groups was retained. Once the final dataset of sequences was determined we removed gap-containing and highly variable columns from the alignment and re-ran MrBayes with mcmc=2,000,000, sump burnin=2000 and sumt burnin=2000 and re-confirmed our inferred tree with maximum likelihood in RAxML.

We also used RAxML to infer a phylogeny of all currently available *D. rerio* and *C. arpio* NLR-B30.2 genes. As described for the conserved NLR genes above we based our phylogeny on an alignment calculated with Clustal-Omega v.1.2, reduced to conserved regions with trimal, and model testing with prottest (JTT+G+F model found to be optimal).

### Divergence analysis

For zebrafish genes in groups 1, 2a, 3a and 3b we calculated all pairwise dNdS values for NACHT domain containing exon and the B30.2 domain independently using the KaKs calculator (D. Wang et al. 2010): We extracted the respective regions from our protein alignment, then used tranalign (Rice, Longden, and Bleasby 2000) to create DNA alignments from these proteins and cds. We then calculated all pairwise comparisons and used paraAT (Zhang et al. 2012) and submitted the resulting alignments to the KaKs calculator independently estimating under the MYN model and the model averaging option with aid of the gnu-parallel tool (Tange 2011). We then used our own iPython (Pérez and Granger 2007) script to sort data and calculated means, medians, and errors in the R statistic software (R-Core-Team 2015).

We also used the tranaligned regions to calculate independent phylogenies for the NACHT and B30.2 exons with RAxML. We loaded the inferred trees into Dendroscope and employed this software to visualise connection between branches belonging to the same gene in both trees.

## ACKNOWLEDGEMENTS

We thank Robert Remy and Giuliano Crispatzu for some early work on this project and Jonathan Wood for verifying genomic locations of assembly components. Financial support was provided by EMBO and the DFG SFB 670 “Zellautonome Immunität” to ML, DFG SFB 680 “Molecular basis of evolutionary innovation” to TW, the HHMI International Early Career Scientist Program (55007424), MINECO (Sev-2012-0208), AGAUR program (2014 SGR 0974), and an ERC Starting Grant (335980_EinME) to FK, the European Molecular Biology Laboratory to JM, the Wellcome Trust to KH (zebrafish genome sequencing project) and the National Human Genome Research Institute (NHGRI) grant HG002659 to GKL (gene annotation), and a grant from the Volkswagen Foundation to PHS. We thank the CHEOPS support team and the Bundesland Nordrhein Westfalen for making HPC applications freely available at the University of Cologne.

## Supplementary Fig. Legends

**SFig 1:**Overview of the entire set of 368 predicted NLR-B30.2 proteins in the zebrafish, based on a Clustal Omega alignment. The original alignment (online supplementary material) was edited by hand to improve the alignments of the N-terminal repeats and the LRR. The colour code for the amino acids was assigned at random. Gaps were introduced in the alignment at the positions of introns (marked by a grey arrowhead below the alignment). Further large gaps are created because some positions are prone to variable and often long insertions or internal duplications (marked by red asterisks below the alignment). The domains are marked below the sequences; domain boundaries within the NACHT domain are entered according to reference (Proell et al. 2008). Positions of introns are marked by grey arrowheads below the sequences.

**SFig 2:**Comparison of FISNA and NACHT domains in groups 1 – 4 in the NLR-B30.2 genes.

Logos generated in Jalview from the alignment in the online supplementary material. Differences in the Walker A motif are diagnostic for groups.

**SFig 3:**Comparison of LRRs in groups 1 – 3 and mouse NLRP3 LRRs.

Logos generated in Jalview from the alignment in online supplementary material.

**SFig 4:**Differences in the B30.2 domain between the indicated groups.

Logos generated in Jalview from the alignment in online supplementary material.

**SFig 5:**Boxplots of dN/dS values for the Fisna and NACHT encoding exons and the exons coding for the B30.3 domain. Median values are indicated by a black bar, means are displayed as a diamond. The y-axis is log-scaled.

**SFig 6:**Phylogenetic maximum likelihood tree of the NLR-B30.2 proteins in *D. rerio* and *C. carpio*.

**SFig 7:**Phylogenetic maximum likelihood tree of the conserved NLR proteins in Metazoa.

